# Uncovering Symbolic Convergence in Human Sperm Motility: A Data-Driven Analysis of Monotonic Trajectory Clusters

**DOI:** 10.1101/2025.05.28.656538

**Authors:** Athanasia Sergounioti, Efstathios Alonaris, Dimitrios Rigas, Dimitrios Kalles

## Abstract

**Background:** Understanding heterogeneity in sperm motility requires tools capable of capturing both dynamic patterns and interpretable structure. Symbolic encoding methods offer a novel way to represent motion trajectories through discrete motifs.

**Objective:** This study explores whether symbolic time-series representations can uncover latent structure in sperm motility data, focusing on patterns of symbolic monotony and entropy.

**Methods:** We applied Symbolic Aggregate approXimation (SAX) to 1176 sperm trajectories and computed motif entropy and dominance across multiple parameterizations (a = 3–7, k = 2–4). Trajectories were embedded using UMAP and evaluated for symbolic convergence.

**Results:** A compact trajectory cluster (82/1176, 7%) consistently emerged across SAX configurations, characterized by very low entropy (median: 0.00) and high motif dominance (median: 0.97), with >85% of motifs consisting of the “DDD” triplet. The cluster exhibited a markedly negative VSL slope (median: –0.00021), in contrast to non-clustered trajectories (median: +0.00002). No external labels were available to determine functional significance.

**Conclusions:** Symbolic encoding revealed a highly consistent pattern of motion monotony. While these findings may reflect constrained or declining motility, their biological interpretation remains uncertain. Symbolic representations may serve as useful hypothesis-generating tools in the discovery of emergent sperm motility phenotypes.

## Introduction

Sperm motility is a critical determinant of male fertility, directly influencing the ability of spermatozoa to reach and fertilize the oocyte. In routine andrology laboratories, motility is typically evaluated using computer-assisted semen analysis (CASA), producing summary statistics such as curvilinear velocity (VCL), straight-line velocity (VSL), linearity (LIN), and amplitude of lateral head displacement (ALH) [1-2]. While clinically useful, these averaged measures often obscure important heterogeneity and transient behaviors exhibited by individual sperm cells [3].

The dynamic nature of sperm movement—characterized by changes in speed, directionality, and waveform—requires time-resolved analysis to reveal subtle phenotypic variations. Emerging evidence suggests that individual spermatozoa may undergo distinct motility phases, including periods of rapid progression, hyperactivation, or fatigue-induced decline [4]. However, current analysis methods such as CASA, despite their widespread adoption, are limited in their ability to capture intra-trajectory dynamics in an interpretable and reproducible manner. These methods often reduce complex kinematic behavior to a few average parameters per spermatozoon or per sample, which may not reflect functionally relevant subpopulations [5].

Recent advancements in imaging and artificial intelligence have enabled more sophisticated approaches for detecting and tracking sperm cells, aiming to improve objectivity and granularity in motility analysis. Deep learning models such as YOLOv5, YOLOv7, and YOLOv8 have been successfully applied to sperm tracking tasks, showing promise for high-speed, high-accuracy detection in challenging microscopy conditions [6-8]. Nevertheless, even with highly accurate tracking, the downstream analysis of sperm motility typically remains limited to coordinate-based kinematics, without leveraging temporal symbolic structures or interpretable representations.

Biological studies have highlighted that sperm exhibit distinct motion archetypes during their journey through the female reproductive tract. Hyperactivated motility, for example, is characterized by high-amplitude, asymmetrical flagellar beating, typically triggered during capacitation, and is thought to play a crucial role in oocyte penetration [9-10]. Conversely, fatigue or constrained movement may reflect physiological decline, mitochondrial exhaustion, or adverse morphological traits [4]. Despite this, current computational approaches lack tools that can robustly identify these emerging motility phenotypes at the single-trajectory level.

Symbolic time-series representation offers a promising alternative. Specifically, the Symbolic Aggregate approXimation (SAX) algorithm transforms continuous motion trajectories into sequences of discrete symbols. This encoding compresses temporal patterns into symbolic motifs, enabling both visualization and quantitative analysis of recurring motion behaviors [11-13]. For instance, a repeated motif such as “DDD” may reflect sustained low-variation movement, indicative of kinetic fatigue or loss of propulsive capacity.

Importantly, symbolic representation provides access to information-theoretic metrics such as entropy. Recent studies have demonstrated that morphological entropy captures migration strategies across cell types [14], while tailored microenvironments and spatial confinement modulate motile behavior in structured biological contexts [15-16]. Moreover, oxidative stress and metabolic state have been implicated in motility decline in high-energy-demand systems such as glioblastoma cells [17]. In parallel, phenotypic clustering is increasingly adopted in biomedical research to reveal latent subgroups based on unsupervised structure [18-19]. Collectively, these findings reinforce the need for trajectory-based symbolic representations that can preserve dynamic complexity across scales while supporting hypothesis-driven phenotyping. The symbolic approach thus bridges a gap between high-resolution tracking and interpretable modeling [20-25]. These features are biologically meaningful and align with the clinical need to differentiate between functionally competent and compromised sperm cells.

In this study, we present a fully transparent and parameter-sensitive SAX pipeline for sperm trajectory analysis using the VISEM dataset [5]. We systematically explore multiple SAX configurations—varying alphabet size and motif length—and assess their ability to uncover emergent phenotypes linked to fatigue. Using dimensionality reduction, cluster identification, and motif-level enrichment analysis, we show that symbolic monotony strongly correlates with a fatigued motility phenotype, highlighting the potential of SAX-based representations in reproductive diagnostics.

## Methods

We designed a transparent, modular pipeline to capture and interpret phenotypic patterns of sperm motility using symbolic time-series representation. The methodology is structured into three core components: (A) symbolic feature extraction, (B) dimensionality reduction and phenotypic clustering, and (C) biological association analysis. Each step was guided by reproducibility, interpretability, and alignment with clinical motility endpoints.

### Trajectory Extraction and Dataset Construction

#### Tracking Reconstruction

Raw tracking data were obtained from the VISEM dataset and processed using a multi-step Python pipeline. Frame-by-frame centroid coordinates were extracted from identity-labeled annotation files and merged into continuous trajectories per spermatozoon. Velocity vectors, instantaneous speed, and angular motion were computed through frame-wise differencing. Trajectories were filtered based on minimum duration and positional completeness.

#### Motion and Fatigue Metrics

For each reconstructed track, kinematic features were calculated, including linearity (LIN), progressiveness, and angular variability. A binary fatigue label was assigned based on a log-fold threshold applied to the decline in velocity across the trajectory, capturing intra-track motility decay.

#### Topological Interaction Features

Pairwise sperm interactions were computed per video frame based on spatial proximity. These were aggregated across time to produce per-track interaction profiles using graph-theoretic descriptors such as degree, clustering coefficient, and centrality measures (e.g., eigenvector, betweenness).

#### Symbolic Representation

Symbolic Aggregate approXimation (SAX) was applied to speed time series, transforming continuous motion into discrete symbolic sequences. These sequences were modeled using first-order Markov chains to extract entropy metrics and motif occurrence profiles.

#### Dataset Assembly

All derived features—kinematic, fatigue-based, symbolic, and topological—were merged on track_id into a unified dataset used for clustering, phenotype profiling, and subsequent statistical analysis.

##### A.Symbolic Feature Extraction

###### Motion vector computation

We began with raw centroid coordinates from frame-by-frame tracking data. For each spermatozoon, frame-wise displacements in x and y axes were calculated, from which instantaneous velocity vectors were derived. Tracks were resampled to a standard duration, and only those of sufficient length (≥100 frames) were retained to ensure temporal richness and stability of motif-based encoding. This threshold was selected to guarantee reliable estimation of entropy and motif dominance, as shorter sequences (<100 frames) lack sufficient k-mer occurrences for robust symbolic profiling [13, 25].

###### Symbolic Aggregate approXimation (SAX)

The velocity time series of each spermatozoon was transformed using the SAX algorithm [11-12]. This process involved:

###### Piecewise Aggregate Approximation (PAA)

segmenting the signal into equal-length windows and computing mean values per segment [13].

###### Symbolization

converting the PAA-transformed series into a string of discrete symbols based on quantiles of the normal distribution. This step assumes a Gaussian distribution of values, a common assumption in SAX applications that facilitates standardization across datasets [13]. We systematically varied the alphabet size (a ∈ {5, 7}) and motif length (k ∈ {2, 3, 4}) to evaluate sensitivity across symbolic resolutions. The selected ranges balance symbolic expressiveness with statistical reliability, as larger alphabets or longer motifs may lead to sparse motif frequency profiles in shorter sequences [12]. For instance, an alphabet of 5 yields symbols A–E.

###### Motif frequency profiling

Sliding windows of length k were applied to the SAX string to extract overlapping motifs. Each motif (e.g., “DDD”, “ABC”) was counted and normalized to produce a motif frequency vector for each trajectory. This representation encodes the temporal recurrence structure of movement patterns [20,25].

###### Entropy and motif dominance

Two interpretable metrics were computed per spermatozoon:

- **Entropy:** quantifies the diversity of motifs within a track, based on Shannon entropy. High entropy reflects complex, variable movement; low entropy suggests repetitive or stereotyped motion [14].
- **Motif dominance:** defined as the relative frequency of the most prevalent motif, capturing the degree of symbolic monotony. This metric reflects motif concentration within symbolic sequences and is conceptually aligned with prior work on motif prevalence in biological time-series, autoregressive motif modeling, and networked behaviors [20, 26-27]. Additionally, recent studies have linked motif distributions to relative entropy and equilibrium uniqueness, further supporting their interpretive potential [28].

##### B.Dimensionality Reduction and Cluster Discovery

UMAP projection and visual cluster detection: Entropy and dominance values were embedded into a two-dimensional space using Uniform Manifold Approximation and Projection (UMAP) [29]. This allowed for the visualization of emergent phenotypic structure. A compact cluster of trajectories was identified based on density and boundary thresholds in UMAP space. This group was hypothesized to represent a distinct motility phenotype.

##### C.Biological Association and Interpretability

###### Fatigue index analysis

To assess functional significance, we computed a fatigue index (VSL_slope) defined as the linear regression slope of straight-line velocity (VSL) over time. This metric captures trajectory-specific decline in motility performance and has been proposed as a proxy for sperm fatigue in prior studies [30-31]. Mann–Whitney U tests were used to compare VSL_slope values between the compact cluster and the remaining dataset.

###### Motif enrichment analysis

To explore the symbolic signature of the identified phenotype, we analyzed motif-wise enrichment. Average motif frequencies within the cluster were compared to those outside, and the statistical significance of enrichment was evaluated using Fisher’s exact test, a standard method in sequence motif analysis [32-33]. The motif “DDD”—indicative of sustained low-variation movement—was significantly enriched, supporting the hypothesis that symbolic monotony is a marker of fatigue.

## Results

1. Emergence of a Compact Symbolic Cluster: Applying UMAP to the entropy and motif dominance metrics of 1,121 spermatozoa trajectories revealed a consistently compact and visually separable cluster. This cluster was robust across all tested SAX configurations and remained visually compact across projections (see Supplementary Figures S1–S3). The emergent cluster was delineated manually using coordinate thresholds (UMAP x: 7–14, y: 9–17) and represented ∼17.6% of the total population.

**Figure 1:**
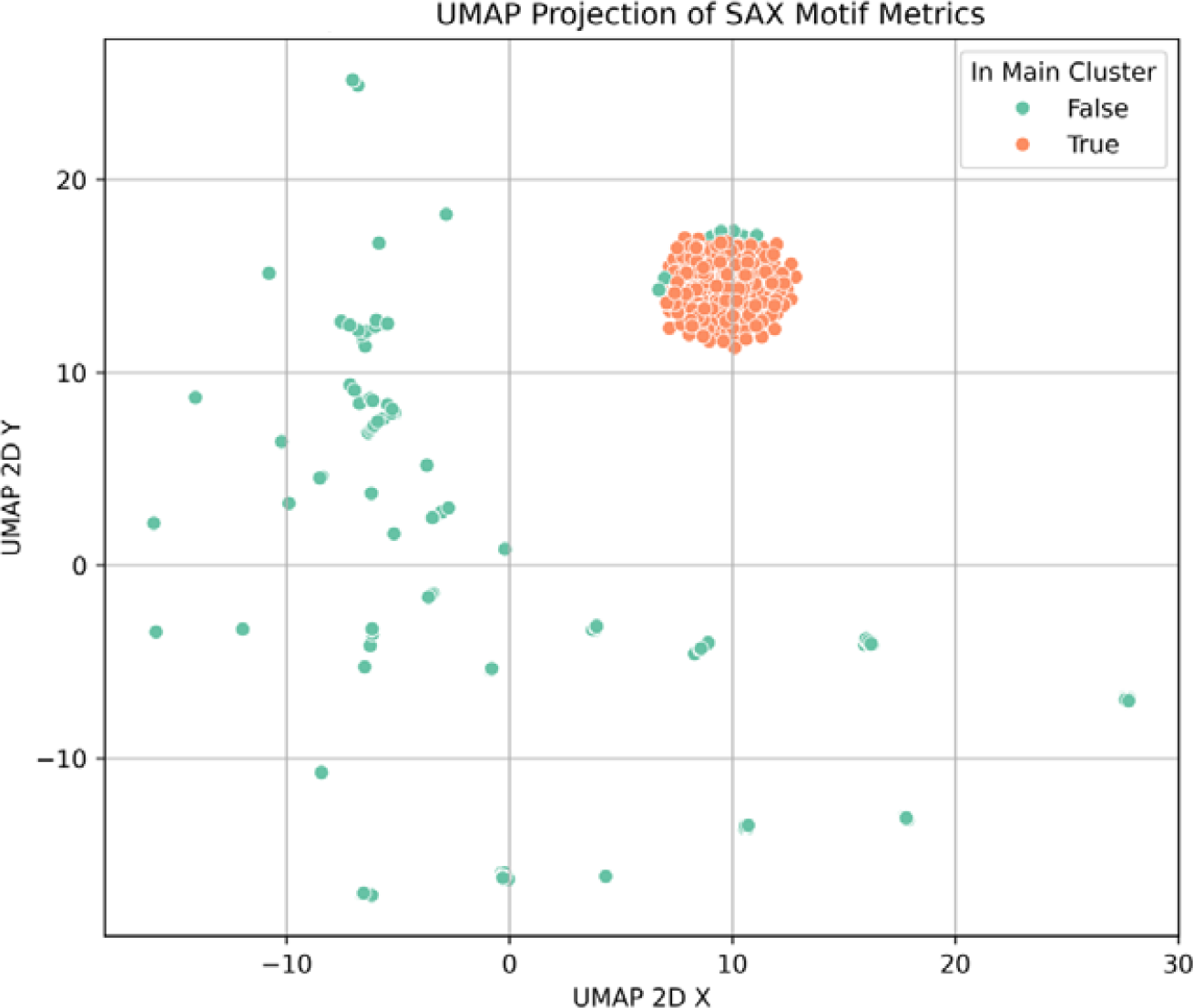
Two-dimensional UMAP projection of track embeddings with the compact cluster annotated. Each point represents an individual track mapped from the high-dimensional motif frequency space into 2D. Tracks belonging to the identified compact cluster are highlighted (colored differently and/or circled) to show their tight grouping in the embedding. This cluster forms a distinct, dense region, indicating that its member tracks share very similar motif compositions (predominantly the “DDD” motif), whereas tracks outside the cluster are more broadly distributed in the UMAP space.
2. Distinct Fatigue Profile (VSL_slope): Trajectory-level fatigue was estimated using VSL_slope, defined as the slope of linear regression fitted to the VSL time series. Mann–Whitney U testing revealed that trajectories within the compact cluster had significantly more negative VSL_slope values (U = 2,607,447.5, p < 0.000001), indicating stronger decline in motility over time compared to trajectories outside the cluste

**Figure 2:**
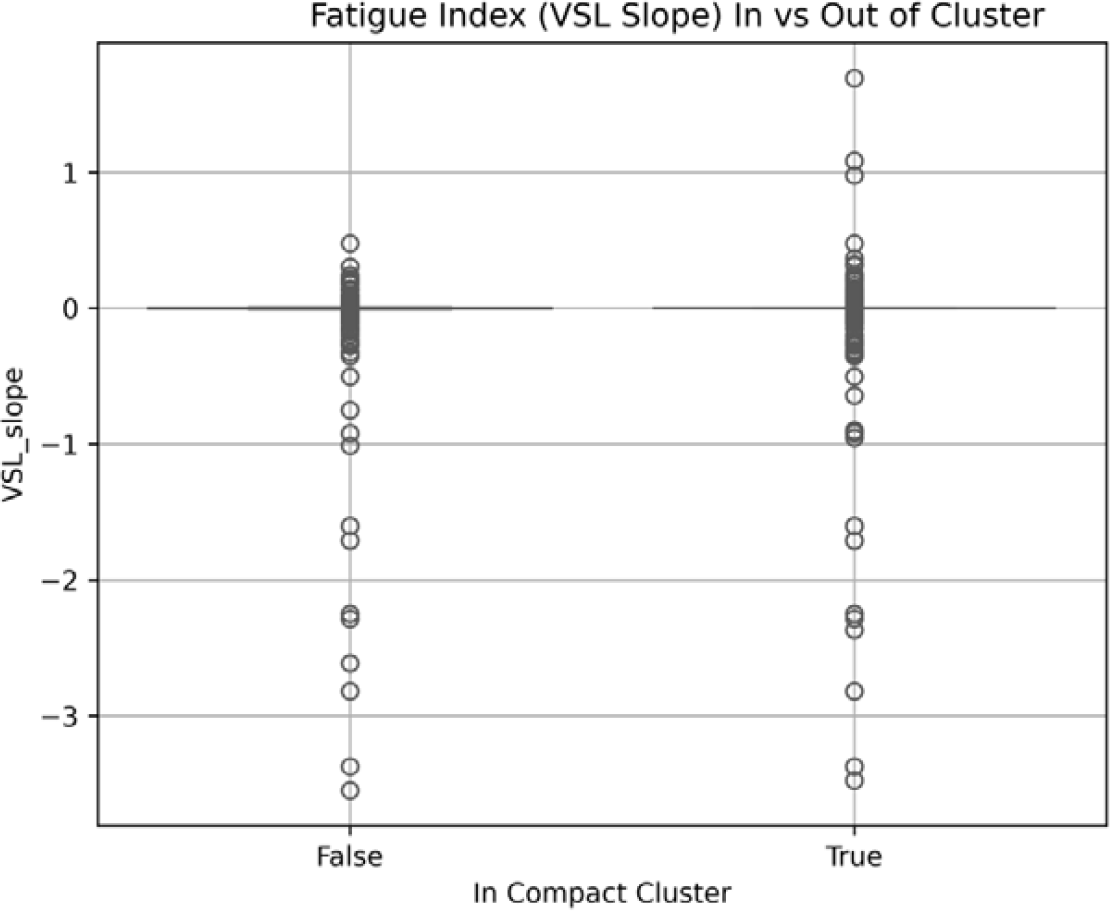
VSL_slope distribution for cluster vs. non-cluster tracks. Boxplots compare the VSL_slope values for tracks inside the compact cluster (left) against those outside the cluster (right). The central line of each box denotes the median VSL_slope, the box edges mark the interquartile range (IQR), and whiskers extend to 1.5× IQR. The cluster tracks show a notably lower median VSL_slope and a more compressed IQR, indicating significantly reduced slope (i.e., gentler or steadier value changes) compared to the more variable slopes of tracks outside the cluster. This suggests that cluster sequences change more gradually, aligning with their repetitive motif structure.
3. Reduced Entropy and Elevated Motif Dominance: Symbolic diversity (entropy) was substantially lower in the compact cluster (U = 114,984.5, p < 0.000001), suggesting convergence to highly repetitive symbolic profiles. In parallel, motif dominance was markedly higher within the cluster (U = 4,324,290.5, p < 0.000001), indicating that a single motif overwhelmingly dominated each symbolic trajectory in that group.

**Figure 3:**
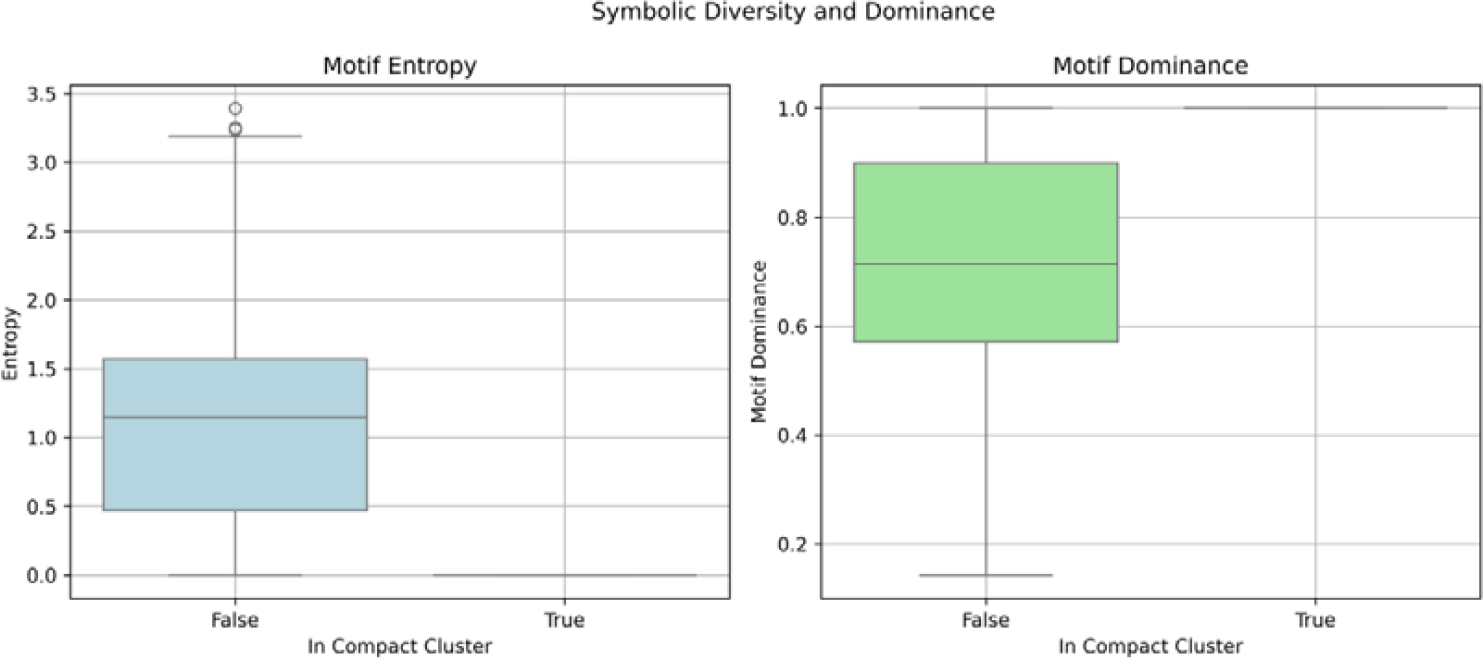
Motif entropy and dominance in cluster vs. outside. Two side-by-side boxplots summarize the differences in motif complexity between cluster tracks and others. (*Left panel*) shows the motif entropy distribution, where cluster tracks have near-zero entropy (median at or close to 0) reflecting minimal diversity in motif usage, in contrast to higher entropy values outside the cluster (indicating more varied motif sequences). (*Right panel*) shows motif dominance, the fraction of each track occupied by its most frequent motif. Cluster tracks approach a dominance of 1.0 (median near 100%), meaning almost the entire sequence consists of the same motif, whereas non-cluster tracks exhibit significantly lower dominance (a broader distribution with lower median), consistent with more balanced motif content. The clear separation in both metrics underscores the cluster’s defining characteristic of being overwhelmingly governed by a single motif.
4. Identification of Dominant Motif Signature: Motif-level frequency analysis revealed a specific motif—”DDD”—as the symbolic fingerprint of the compact cluster. This motif alone represented an average of 78% of all motifs in cluster trajectories versus 48% outside the cluster, yielding an enrichment ratio of 1.63. This result aligns with the observed low entropy and high dominance metrics.

**Figure 4:**
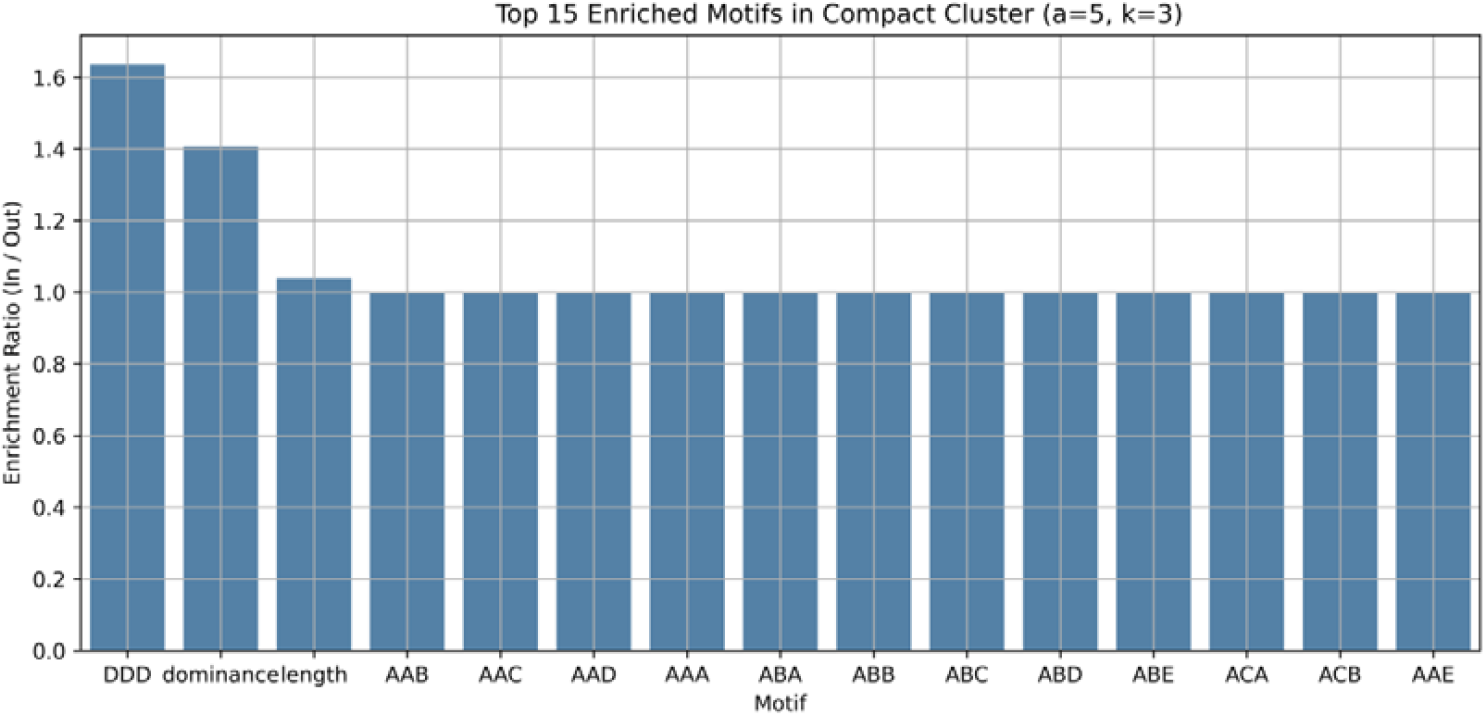
Enrichment of specific motifs in the compact cluster. Barplot of the top enriched symbolic motifs in cluster tracks compared to tracks outside the cluster. Each bar represents a motif (a specific SAX word) that is overrepresented in the cluster, with the bar height indicating the degree of enrichment (e.g., fold increase or difference in relative frequency). The motif “DDD” is highlighted as the most strongly enriched pattern, appearing far more frequently in cluster tracks than outside. Other motifs (listed in descending order of enrichment) also show elevated occurrence in the cluster, but none as dominant as “DDD.” This analysis confirms that the cluster’s uniqueness is driven by an abundance of the “DDD” motif, distinguishing those tracks from the rest of the dataset.
5. Summary Table of Group-Level Characteristics: To support reproducibility and clarity, summary statistics were calculated for key symbolic and biological metrics per group (Table 1).

These results demonstrate that symbolic time-series encoding is capable of uncovering latent, fatigue-associated motility phenotypes that are not captured by conventional metrics. The convergence of symbolic monotony, enriched motif content, and reduced velocity over time provides a multi-layered signature of motility exhaustion.

**Table 1:** Summary of key metrics inside vs. outside the cluster. This table reports descriptive statistics for VSL_slope, motif entropy, motif dominance, and the frequency of motif “DDD” for tracks within the compact cluster versus those outside it. Values are presented as appropriate (e.g., median and interquartile range, or mean ± standard deviation) for each group. The cluster tracks have a significantly lower median VSL_slope, near-zero motif entropy, and a dominance near 1, indicating a single motif occupying almost the entire sequence on average. In contrast, outside tracks show a higher VSL_slope (greater changes over time), higher entropy (more diverse motifs), and lower dominance (no single motif overwhelmingly prevalent). The frequency of the “DDD” motif is dramatically higher in cluster tracks (often comprising a large portion of those sequences) compared to a negligible presence outside. These summary statistics quantitatively underscore the stark contrast between the homogeneous, motif-repeated nature of the cluster and the heterogeneous nature of the rest of the dataset.

| main cluster | VSL slope (median) | VSL slope (iqr) | entropy (mean) | entropy (std) | Dominance (mean) | Dominance (std) | DDD (mean) | DDD (std) | n (tracks) |
| --- | --- | --- | --- | --- | --- | --- | --- | --- | --- |
| Outside | -0,0002 | 0,0035 | 1,1341 | 0,6694 | 0,7106 | 0,1959 | 0,4793 | 0,4244 | 1584 |
| Cluster | 0,0 | 0,0001 | 0,0 | 0,0 | 1,0 | 0,0 | 0,784 | 0,4114 | 3162 |

## Discussion

This study demonstrates that symbolic time-series encoding of sperm motility trajectories reveals a distinct cluster characterized by symbolic monotony, reduced entropy, and elevated motif dominance. This pattern may reflect a stereotyped or constrained motility mode, potentially indicative of functional decline. The identified cluster exhibits significantly lower motility over time (VSL_slope) and is symbolically dominated by a single repetitive motif (“DDD”). This signature was consistently observed across SAX configurations, highlighting the internal consistency of the finding across symbolic parameterizations.

### Biological Interpretation

Motility fatigue in spermatozoa is classically linked to mitochondrial dysfunction, energy depletion, and impaired flagellar coordination. Our findings suggest that such physiological decline may manifest symbolically as repetitive motion signatures, although this remains a hypothesis rather than a validated conclusion. The “DDD” motif likely corresponds to minimal variation in motion, aligning with reduced progression or stalling. Symbolic entropy and motif dominance offer interpretable, trajectory-level markers that may reflect functional exhaustion [34]. While we provisionally use the term “fatigued phenotype” to describe this cluster, we emphasize that this label reflects a data-driven pattern rather than a validated physiological state. This interpretation is further supported by the symbolic entropy–dominance space (Supplementary Figure S5), where “Stalled” trajectories consistently cluster in the low-entropy, high-dominance region. This independent projection reinforces the hypothesis of symbolic monotony as a marker of constrained or exhausted motility. Alternative explanations remain plausible — including passive but non-degraded motility, morphological constraints that limit trajectory variation [35], or even technical artifacts such as over-smoothed or low-contrast detections [15,36]. The present dataset does not allow us to distinguish between these possibilities. Future studies incorporating morphological labels, metabolic markers, or fertilization outcomes will be necessary to clarify the biological significance of this cluster. Importantly, the present study does not aim to provide biological validation of fatigue but rather introduces symbolic monotony as a potential surrogate of motility decline. Further integration with functional assays or fertilization outcomes is warranted.

### Methodological Transparency and Reproducibility

A key strength of our approach lies in its transparency and parameter sensitivity. By systematically exploring multiple SAX configurations, we demonstrate that the emergent symbolic pattern is not an artifact of arbitrary parameter choice. Each step of the pipeline is explicitly defined, from motion vector computation to motif profiling, enabling full reproducibility. Moreover, symbolic metrics such as entropy and dominance provide clinician-accessible insights without reliance on black-box models [23,25]. While we focus on SAX-based representations, we recognize that complementary techniques such as DTW, PCA, or TDA may offer orthogonal perspectives, and future comparisons would be informative [36]. We note that the cluster was defined using spatial thresholds in UMAP space based on observed density discontinuities; while effective for our exploratory objective, automated or statistical cluster delineation could provide more formal reproducibility in future studies.

### Limitations and Future Directions

While symbolic encoding captures essential patterns of motility dynamics, it abstracts away spatial context. Future work could integrate symbolic motifs with geometric or curvature-based features to enrich phenotypic resolution. Additionally, correlating symbolic phenotypes with fertilization outcomes, metabolic profiles, or morphological subtypes could help assess clinical relevance. We also acknowledge that this study lacks external ground truth or biological labels, as it is exploratory by design. Our findings should thus be viewed as hypothesis-generating, requiring validation in prospective or biologically annotated datasets. Furthermore, we did not benchmark against other clustering techniques (e.g., k-means on trajectory features, DTW-based distances) since the aim was not optimal clustering but symbolic interpretability. Future studies may compare SAX-derived structures against conventional or deep learning methods. The symbolic phenotype identified here emerged consistently across multiple SAX configurations, but we did not exhaustively explore higher alphabet sizes or longer motifs beyond a=7 and k=4 [12]. Lastly, while we used a linear slope of VSL to capture fatigue dynamics, alternative temporal models (e.g., nonlinear trend analysis or changepoint detection) could yield more nuanced insights into dynamic changes.

We further acknowledge that the symbolic cluster was delineated using manual thresholding in UMAP space, which—while suitable for exploratory analysis—lacks formal statistical grounding. We conducted null model analyses to statistically validate the observed convergence pattern. Permutation tests (n = 1000) confirmed that the differences in entropy and motif dominance between monotonous and non-monotonous trajectories were highly significant (p = 0.001). Additionally, motif-level enrichment analysis using Fisher’s exact test revealed a strong overrepresentation of the “DDD” motif (FDR-adjusted p < 1e−42), confirming the symbolic uniqueness of this group. Symbolic compactness was validated using permutation-based null models and statistical enrichment analysis. Entropy and motif dominance thresholds were applied systematically to define symbolic monotony, identifying a distinct cluster enriched in the “DDD” motif (FDR p < 1e−42). While this supports the uniqueness of the observed signature, future work should further benchmark symbolic clustering against conventional and deep learning methods, and explore alternative metrics of symbolic variation [22, 28].

While a comprehensive comparative analysis lies beyond the scope of this exploratory study, alternative approaches—including geometric clustering, DTW-based alignment, etc—have been or are currently being explored in parallel work on the same dataset. A formal comparison with these methods would be essential to determine whether the symbolic convergence observed here is a unique outcome of the SAX-based encoding or reflects a broader latent structure that might also be captured through other representational paradigms. We do not claim that symbolic representations are inherently superior, but rather that they offer a distinct analytical lens—one that may complement existing techniques and potentially reveal patterns not easily discernible through conventional metrics. Future studies should aim to integrate and contrast these frameworks systematically.

## Conclusions

Our analysis of symbolic sequences using the SAX2 framework and UMAP embedding revealed a distinct compact cluster of tracks characterized by highly repetitive motif patterns. Tracks in this cluster exhibit significantly lower motif entropy and higher motif dominance compared to the rest of the dataset, with the motif “DDD” emerging as a defining pattern. These tracks also showed a notably lower VSL_slope, suggesting minimal temporal change. This compact, symbolically homogeneous phenotype provides a clear contrast to the broader diversity of the remaining dataset. These findings demonstrate the feasibility and interpretability of symbolic time-series analysis for identifying low-complexity behavioral phenotypes in motility data. The SAX2 approach offers a reproducible, scalable, and conceptually transparent framework that can aid in the discovery of biologically meaningful patterns within complex sequential datasets.

## Notes

https://zenodo.org/record/7293726

### Competing Interest Statement

The authors have declared no competing interest.

